# Exopolysaccharide biosynthetic glycoside hydrolases can be utilized to disrupt and prevent *Pseudomonas aeruginosa* biofilms

**DOI:** 10.1101/032714

**Authors:** Perrin Baker, Preston J. Hill, Brendan D. Snarr, Noor Alnabelseya, Mathew J. Pestrak, Mark J. Lee, Laura K. Jennings, John Tam, Roman Melnyk, Matthew R. Parsek, Donald C. Sheppard, Daniel J. Wozniak, P. Lynne Howell

## Abstract

Bacterial biofilms are a significant medical challenge as they are recalcitrant to current therapeutic regimes. A key component of biofilm formation in the opportunistic human pathogen *Pseudomonas aeruginosa* is the biosynthesis of the exopolysaccharides Pel and Psl, which are involved in the formation and maintenance of the structural biofilm scaffold and protection against antimicrobials and host defenses. Given that the glycoside hydrolases – PelA_h_ and PslG_h_ – encoded in the *pel* and *psl* biosynthetic operons, respectively, are utilized for *in vivo* exopolysaccharide processing, we reasoned that these would provide specificity to target *P. aeruginosa* biofilms. Evaluating these enzymes as potential therapeutics, we demonstrate that these glycoside hydrolases selectively target and degrade the exopolysaccharide component of the biofilm matrix and that nanomolar concentrations of these enzymes can both prevent biofilm formation as well as rapidly disrupt preexisting biofilms *in vitro*. This treatment was effective against clinical and environmental *P. aeruginosa* isolates and reduced biofilm biomass by 58–94%. These non-cytotoxic enzymes potentiated antibiotics as the addition of either enzyme to a sub-lethal concentration of colistin reduced viable bacterial counts by 2.5 orders of magnitude. Additionally, PelA_h_ was able to increase neutrophil killing by ~50%. This work illustrates the feasibility and benefits of using bacterial exopolysaccharide biosynthetic glycoside hydrolases and synthetic biology to develop novel anti-biofilm therapeutics.

## Introduction

Bacterial biofilms provide a protective lifestyle for bacteria and are extremely challenging and costly to treat, as they are notoriously recalcitrant to antibiotics and host defenses (*1–5*). It is estimated that 65–80% of all human bacterial infections are biofilm related (*6*). Biofilms are complex communities of bacteria embedded in a extracellular matrix composed of proteins, extracellular DNA (eDNA) and exopolysaccharides. The exopolysaccharide component of biofilms can function to impair antibiotic penetration (*7, 8*) and provide a barrier against phagocytosis by host immune cells (*9*). Given the rise of antibiotic resistance and the discovery that sub-inhibitory concentrations of antibiotics and antimicrobial compounds can promote biofilm formation, there is an urgent need for novel and effective treatments that target and disrupt biofilms (*10–12*). While several compounds have been demonstrated to inhibit *Pseudomonas aeruginosa* biofilm formation (*13–22*), most are unable to disrupt established biofilms, which is more clinically relevant.

*P. aeruginosa* is a ubiquitous, Gram-negative, opportunistic pathogen that is commonly associated with nosocomial infections (*23*). Mortality associated with *P. aeruginosa* infections is high (*24*), and the emergence of multi- and even pan-resistance to antimicrobials have been reported (*25*). The bacterium has the genetic capacity to synthesize at least three different biofilm exopolysaccharides: Psl, Pel and alginate. These polysaccharides are integral components of the extracellular biofilm matrix (*26, 27*). Although alginate production results in a mucoid phenotype and is correlated with chronic infection and poor prognosis in patients with cystic fibrosis, this exopolysaccharide is dispensable for biofilm formation in non-mucoid *P. aeruginosa* strains (*28–30*). Psl is a neutral polysaccharide composed of a pentasaccharide repeat unit of D-mannose, L-rhamnose and D-glucose (*31*) while Pel has been recently identified as a cationic polysaccharide composed of partially acetylated *N*-acetyl-D-glucosamine and *N*-acetyl-D-galactosamine (*32, 33*). Psl, and under some circumstances, Pel, function to facilitate initial surface attachment (*31, 32, 34–36*). Both exopolysaccharides play a significant role in the formation and maintenance of the biofilm architecture (*7, 37, 38*)*. P. aeruginosa* strains with genetic deletions of the *pel* and *psl* operons are profoundly impaired in biofilm formation and virulence in a mouse model of acute infection (*29*). Psl provides protection against neutrophil phagocytosis and antibiotics with diverse biochemical properties (*8, 9*) while Pel enhances resistance to aminoglycosides (*7, 39*). While the preference of Pel or Psl is often strain specific, many isolates are capable of switching between the synthesis of Pel and Psl in response to stress to maintain infection in the host (*40, 41*). This adaptive mechanism underscores the importance of developing therapies that target both exopolysaccharides.

The utilization of Pel and Psl for antibiotic resistance and virulence makes them attractive targets for the prevention and treatment of *P. aeruginosa* infection. However, compounds that target and disrupt these integral biofilm components have not been identified. We therefore sought to identify enzymes that selectively target and degrade Psl and Pel. One common feature shared among many exopolysaccharide biosynthetic operons is the presence of a gene encoding a glycoside hydrolase that is proposed to hydrolyze the exopolysaccharide produced by the biosynthetic pathway (*33, 42–46*). We have exploited these naturally derived glycoside hydrolases as a method of biofilm prevention and dispersal. We demonstrate that the addition of low nanomolar concentrations of these enzymes – PelA_h_ and PslG_h_, – can both prevent biofilm formation as well as disrupt existing biofilms of laboratory, clinical and environmental isolates of *P. aeruginosa in vitro* at nanomolar concentrations. In addition to disrupting biofilms, these non-cytotoxic enzymes potentiate antibiotics and enhance susceptibility to killing by neutrophils. These studies provide us with a method to find enzymes with anti-biofilm activity for the treatment and eradication of chronic bacterial infections.

## Results

**Design and Expression of Enzymatically Active Glycoside Hydrolases**. The Gramnegative bacterium *Actinobacillus actinomycetemcomitans* secretes a biofilm-modifying glycoside hydrolase that allows for the dispersal from the biofilm (*47*), while the Grampositive *Bacillus licheniformis* secrets nucleases to disrupt established biofilms and inhibit biofilm formation of competing bacteria (*48*). To date, no secreted biofilm dispersal enzymes have been identified in *P. aeruginosa*. However, previous bioinformatics analyses have identified PslG and the N-terminal domain of PelA as putative periplasmic glycoside hydrolases encoded in the *psl* and *pel* biosynthetic operons, respectively (*33*). We recently purified and functionally characterized PslG_31–442_, a member of glycoside hydrolase family 39 herein referred to as PslG_h_. This construct removes an N-terminal transmembrane domain, producing a soluble, catalytically active, glycoside hydrolase domain that can hydrolyze Psl (*49*). PelA is a bifunctional protein that contains at least two catalytic domains, a putative glycoside hydrolase domain and a CE4 deacetylase domain (*33*). Based on bioinformatics prediction with the CAZymes Analysis Toolkit (*50*), we generated a PelA_47–303_ construct, herein referred to as PelA_h_, to explicitly study the activity of the glycoside hydrolase domain. This construct was soluble and could be purified to homogeneity using Ni-NTA purification and size-exclusion chromatography, yielding 50 mg of protein per liter of bacterial cell culture.

**Enzyme-Catalyzed Disruption of Biofilms**. We hypothesized that the exogenous application of the glycoside hydrolases PelA_h_ and PslG_h_ to Pel and Psl dependent biofilms, respectively, would result hydrolysis of the exopolysaccharides, thereby disrupting these established biofilms. To assay for biofilm disruption, biofilms were produced using the following strains: PA14 (Pel dependent matrix), PAO1 (Psl dependent matrix), and L-arabinose inducible *P. aeruginosa* PAO1 Δ*wspF* Δ*psl P*_BAD_*pel*, and PAO1 Δ*pelF P*_BAD_*psl*, which exclusively produce Pel and Psl, respectively. PelA_h_, and a putative catalytically inactive E218A variant (PelA_h_ E218A) were applied to Pel-dependent biofilms, while PslG_h_ and an inactive E165Q/E276Q variant (PslG_h_ E165Q/E276Q) were applied to Psl-dependent biofilms. Confocal microscopy coupled with fluorescently-labeled lectin staining with lectins specific for Psl (*Hippeastrum hybrid*, HHA) and Pel (*Wisteria Floribunda*, WFL) demonstrated that catalytically active hydrolases, but not the inactive variants, were capable of degrading the Pel-dependent and Psl-dependent biofilm biomass based on the elimination of fluorescent signal following treatment (**Fig 1**).

**Figure 1.**
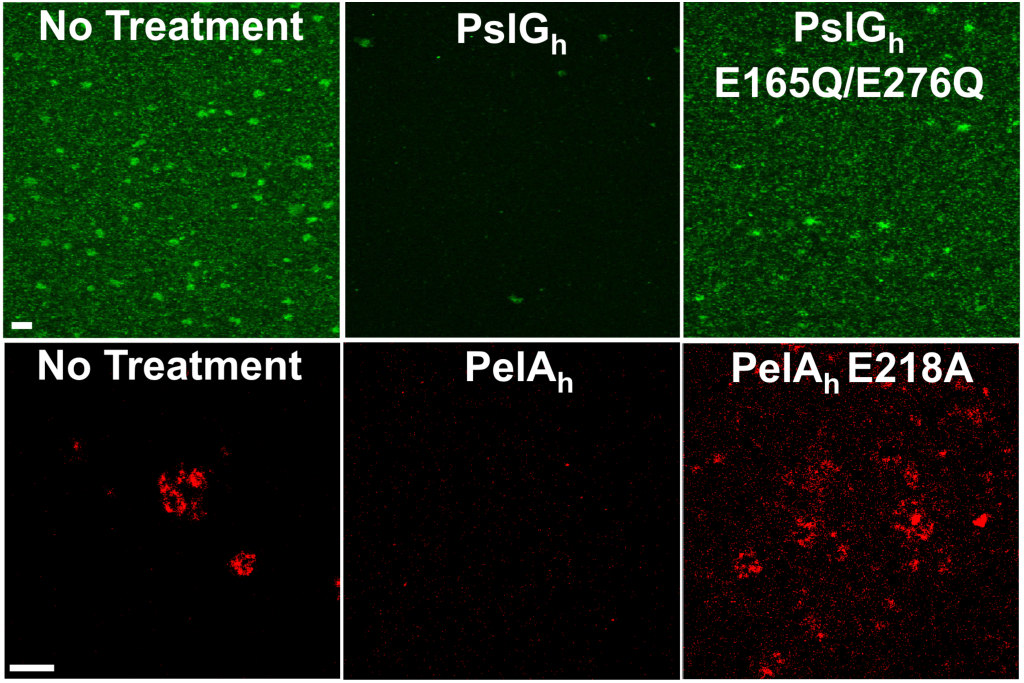
The glycoside hydrolases PslG_h_ and PelA_h_ hydrolyze the exopolysaccharides Pel and Psl in a biofilm. Representative confocal images of Psl biofilms grown statically for 24 h (Top) and Pel biofilms cultivated for 48 h (Bottom) under flow conditions and treated with the indicated hydrolases. Biofilms were stained with the HHA Psl-specific lectin (green) and WFL Pelspecific lectin (red) (Scale bars: 30 μm).

Crystal violet staining was subsequently utilized to quantify the effect of hydrolase treatment on the total biofilm biomass. A 2 h treatment of a Pel-dependent biofilm with PelA_h_ resulted in disruption of 99% of the biomass, while both the PelA_h_ E218A variant and PslG_h_, added in 100-fold excess compared to PelA_h_, exhibited no significant difference compared to that of the untreated biofilm (**Fig 2A**). Similar results were obtained for Psl-dependent biofilms wherein only treatment with a catalytically active PslG_h_ resulted in a 98.5% reduction in biofilm biomass. The activity of both enzymes was observed to be dose dependent. When incubated with established biofilms for 1 h, PelA_h_ exhibited an EC_50_ of 35.7 ± 1.1 nM, while PslG_h_ had an EC_50_ of 12.9 ± 1.1 nM (**Fig 2B**). Time course experiments using fixed concentrations of PelA_h_ and PslG_h_ revealed a continuous decrease in biofilm biomass over time (**Fig S1**). Combined, these results indicate that biofilm-dispersal is catalytic, rapid, and exopolysaccharide specific.

**Figure 2.**
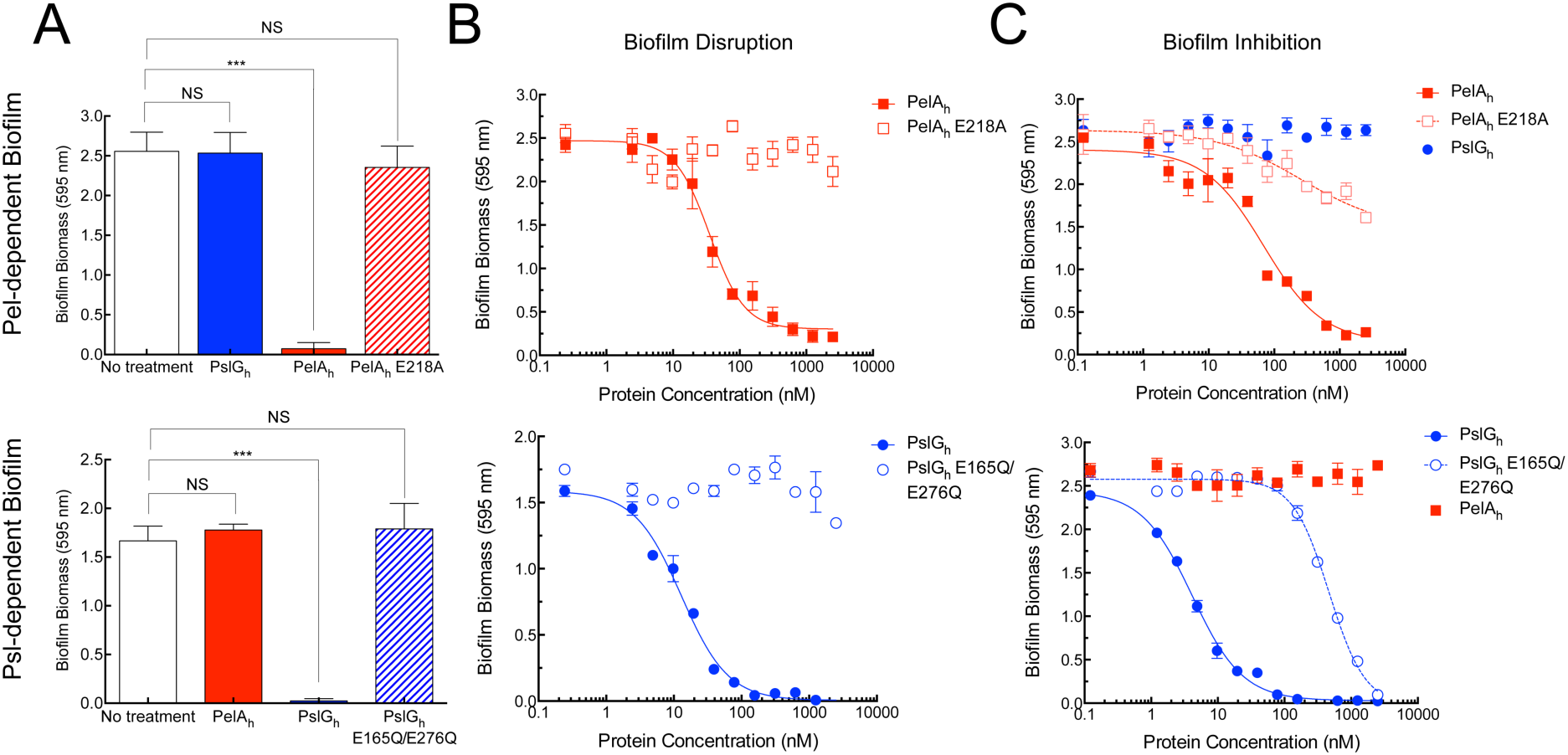
*P. aeruginosa* biofilm inhibition and disruption by glycoside hydrolases (A) Crystal violet staining of biofilms following the exogenous addition of glycoside hydrolases or catalytic variants. (B) Dose-response curves to examine the dispersal of biofilm biomass by the exogenous treatment of each glycoside hydrolase and variant. (C) Dose-response curves to examine the prevention of biofilm biomass in the presence of various glycoside hydrolases. Each data point represents the mean from three independent experiments of *n* = 3 crystal violet microtiter plate wells. Error bars indicate SEM. The *** indicates a significant difference (p ≤ 0.001) and NS represents no significant difference.

**Glycoside Hydrolases Inhibit Biofilm Formation but not Bacterial Growth**. Since the glycoside hydrolases were effective at disrupting established biofilms, we next sought to determine whether the application of enzyme to bacterial culture could be utilized as a prophylactic strategy to prevent biofilm formation. The addition of PelA_h_, but not PslG_h_, to Pel-producing *P. aeruginosa* abrogated biofilm formation. A dose-titration indicated that Pel-biofilms could be prevented over 24 h by PelA_h_ addition with an EC_50_ of 69.3 ± 1.2 nM (**Fig 2C**). As visualized in borosilicate tubes, PelA_h_ prevented pellicle biofilm at the air-liquid interface and bacterial cells grew exclusively in the planktonic state (**Fig S2**). Interestingly, addition of PelA_h_ E218A, which cannot catalyze the dispersal of biofilms, resulted in a statistically significant reduction in biofilm biomass at concentrations ≥500 nM. While an accurate EC_50_ value could not be readily determined for this catalytic variant, 5 μM of the enzyme variant (>70 times greater than the EC_50_ of the wild-type) resulted in <50% reduction of the biomass relative to untreated cells. The presence of ≥10 μM PslG_h_ did not affect the ability of *P. aeruginosa* to form Pel-dependent biofilms. Consistent with results for PelA_h_ on Pel-dependent biofilms, the addition of 1 μM PslG_h_ to Psl producing cultures under biofilm forming conditions resulted in a complete inhibition of biofilm formation, and a dose-titration indicated that PslG_h_ had an EC_50_ of 4.1 ± 1.1 nM over 24 h. To examine whether the effect was the direct result of PslG_h_ activity, the catalytically inactive variant PslG_h_ E165Q/E276Q was tested. This variant was >100-fold less effective at biofilm prevention (EC_50_ of 466.5 ± 1.1 nM) relative to the catalytically active enzyme. Addition of ≥10 μM PelA_h_, had no affect on Psl-dependent biofilm production.

We next examined the length of time that the glycoside hydrolases could prevent biofilm formation. A single dose of PelA_h_ or PslG_h_ prevented biofilm formation for 48 and 72 h, respectively. The formation of a Pel-dependent biofilm at 72 h was associated with proteolytic degradation of PelA_h_, while no degradation of PslG_h_ was observed over the entirety of the experiment (**Fig S3**). The growth rate of *P. aeruginosa* PAO1 exposed to ≥20 μM of either glycoside hydrolase was unaffected over six hours of static growth when compared to a no treatment control (**Fig S4**). The absence of enzyme cross-reactivity between exopolysaccharides indicates that biofilm inhibition is highly specific. This result, combined with bacterial growth curves, demonstrates that exogenous glycoside hydrolases do not impede biofilm formation by altering cell viability and growth.

**Biofilm-Dispersing Enzymes are Non-Cytotoxic**. Since exogenous PelA_h_ and PslG_h_ did not affect *P. aeruginosa* cell viability and growth, we next sought to examine whether enzyme treatment impacted mammalian cells. IMR-90 human lung fibroblast cells treated for 5 h with concentrations up to 1 mg/mL of either enzyme, ~100-fold above the concentration required for effective biofilm disruption, resulted in no significant difference in cell area or length-to-width ratio (**Fig 3A**). Following 48 h incubation, no significant difference in cellular viability was observed in PelA_h_ and PslG_h_ treated cells regardless of the media used (**Fig 3B**). *Clostridium difficile* toxin TcdB (*51*) was used as a control for cell morphology as it results in cell rounding, while digitonin, which permeabilizes the cells, was used to monitor cellular viability (**Fig 3B**). Western blotting of the media confirmed that PelA_h_ and PslG_h_ remained intact for the 48 h duration of the experiment (**Fig S5**). Together, these results suggest that the enzymes do not interfere with mammalian cell morphology and viability.

**Figure 3.**
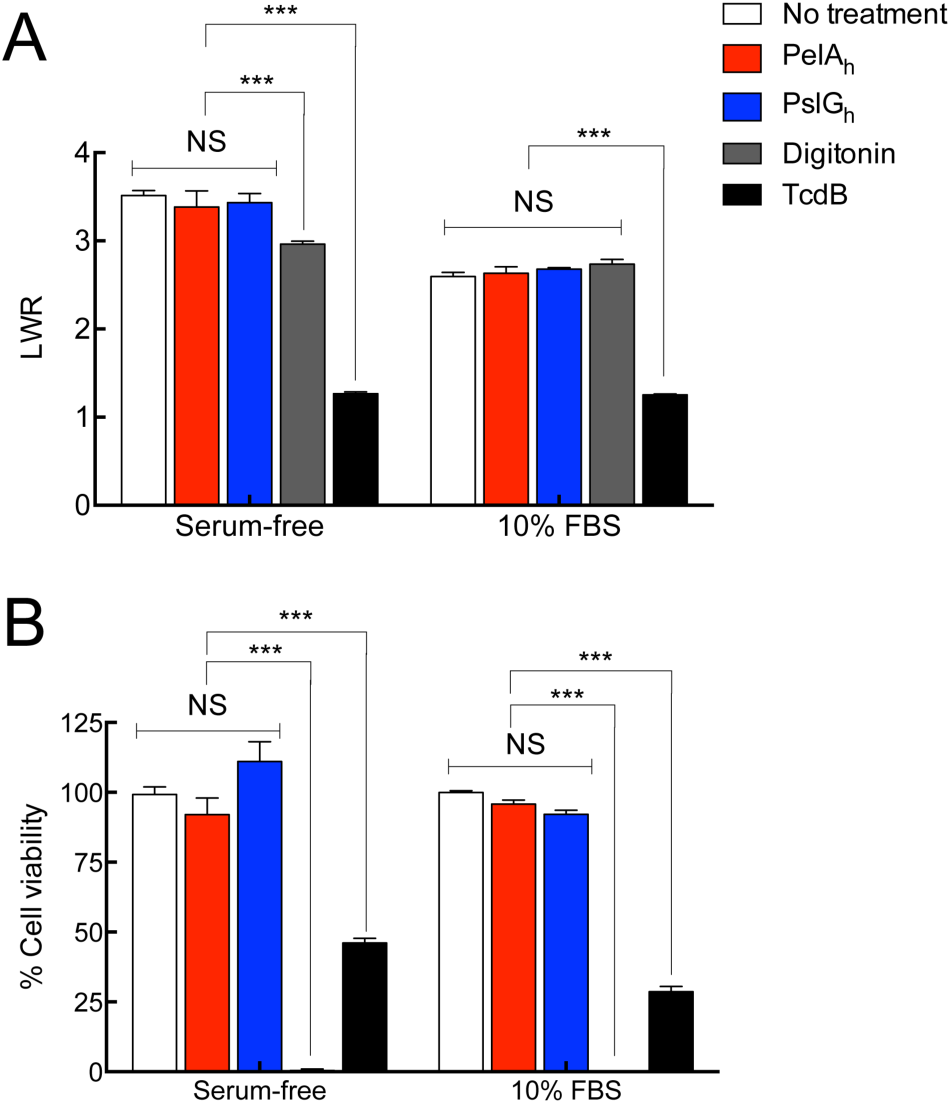
The glycoside hydrolases PelA_h_ and PslG_h_ are non-cytotoxic. (A) IMR-90 cellomics assay to measure the length-to-width ratio (LWR) of the cells using CellTracker Orange CMRA. (B) IMR-90 fibroblast cell viability assay using PrestoBlue^®^ reagent. All data was normalized to a no treatment control (100%). The *C. difficle* toxin TcdB was used as a positive control in cell morphology assays and the detergent digitonin was utilized as a negative control in cell viability assays. Each data point represents the mean from three independent experiments of *n* = 3 from cellomic and PrestoBlue^®^ measurements in microtiter plate well. Error bars indicate SEM. The *** indicates a significant difference (p ≤ 0.001) and NS represents no significant difference.

**Enzymes Potentiate Antibiotics and Ameliorate Human Neutrophil Killing**. Previous studies have demonstrated that both Pel and Psl enhance antibiotic resistance (*7, 8*). We therefore theorized that glycoside hydrolase degradation of these polymers could potentiate the activity of antimicrobial agents. As the antibiotic colistin targets the cell membrane of *P. aeruginosa*, it is active against both the metabolically active and dormant cells found within biofilms (*52*). We therefore examined the effect of combining glycoside hydrolase treatment with sub-inhibitory concentrations of colistin. As predicted, treatment of *P. aeruginosa* Pel- and Psl-dependent biofilm cultures with colistin (50 μg/mL), PelA_h_, or PslG_h_ (2 μM) alone had no effect on the viability of the bacteria (**Fig 4A**). However, the combination of PelA_h_ or PslG_h_ with colistin resulted in a ~2.5 log reduction in bacterial colony forming units. This result indicates that these glycoside hydrolases are compatible with antibiotics and can potentiate the antimicrobial activity of colistin.

**Figure 4.**
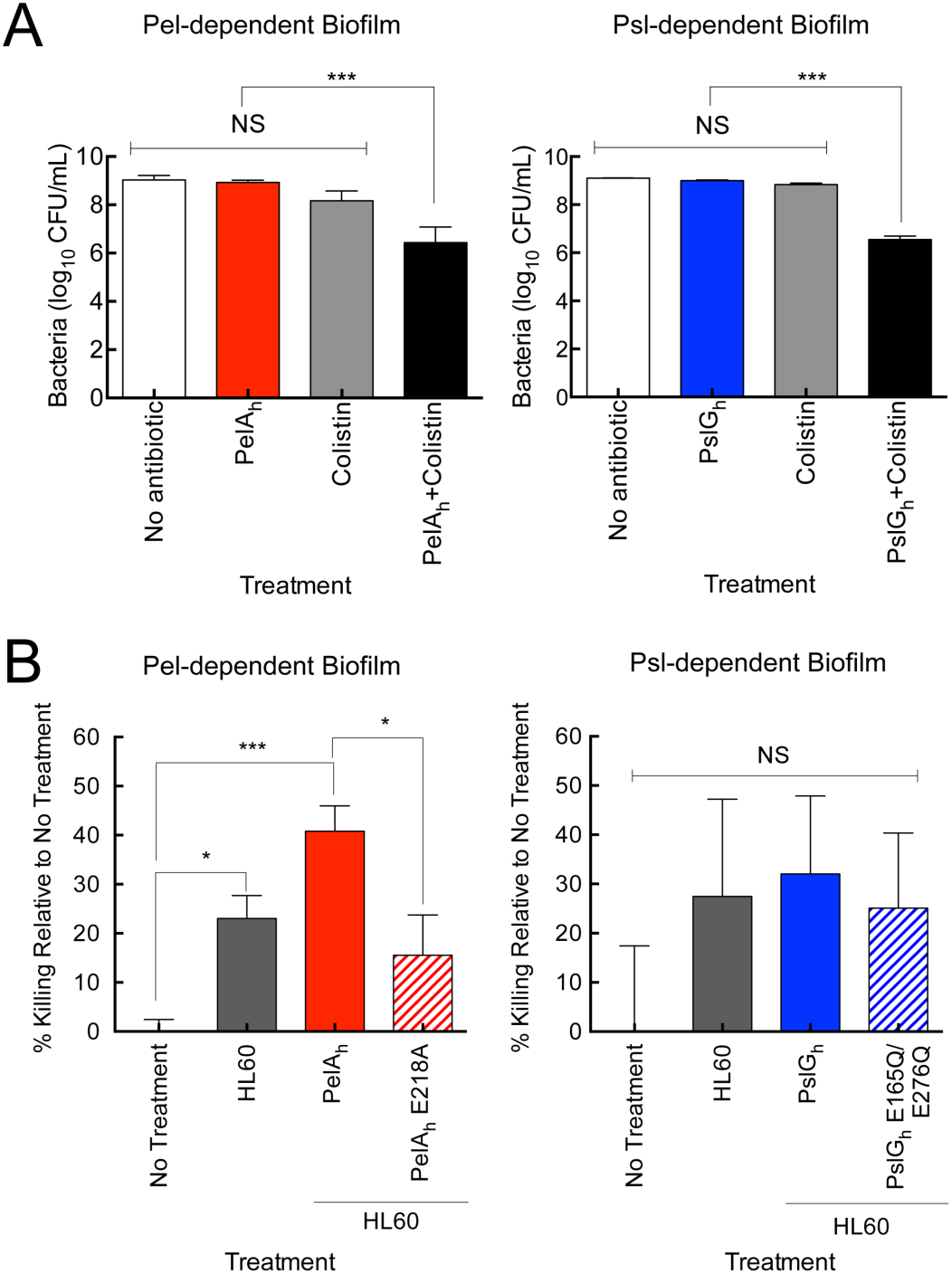
Glycoside hydrolases potentiate antibiotics and increase human neutrophil killing. (A) Cell number count for PAO1 Δ*wspF* Δ*psl* P_BAD_ *pel* (Left) and PAO1 Δ*pelF* P_BAD_ *psl* (Right) and in the presence of glycoside hydrolases, antibiotic and a combined enzyme + antibiotic treatment. Mean was calculated from LB agar plate counts from 3 independent experiments. (B) HL60 neutrophil killing of strain PAO1 Δ*wspF* Δ*psl* P_BAD_ *pel* and PAO1 Δ*pelF* P_BAD_ *psl* following biofilm formation and treatment with PelA_h_ and PslG_h_ and their catalytic variants, respectively. Percent killing was normalized to a no treatment control. Error bars indicate SEM. The * indicates a significant difference of (p ≤ 0.05), *** indicates (p ≤ 0.001) and NS represents no significant difference.

Since *P. aeruginosa* exopolysaccharide enhance resistance to human neutrophil killing, we next investigated whether glycoside hydrolase treatment could enhance susceptibility to immune-mediated killing. It has been demonstrated *in vitro* that exopolysaccharide produced by *P. aeruginosa* enhance resistance to neutrophils by inhibiting efficient opsonization and reducing reactive oxygen species production (*9*). To determine if hydrolase treatment could impact neutrophil killing, the ability of PelA_h_ and PslG_h_ to enhance the susceptibility of *P. aeruginosa* to the human HL-60 neutrophil cell line was examined. Treatment of Pel-containing *P. aeruginosa* biofilms with PelA_h_ increased the degree of HL-60-mediated microbial killing from approximately 22% to 42% (**Fig 4B**). This was not observed in a PelA_h_ E218A variant indicating that the enhanced susceptibility to neutrophils is due to the catalytic activity of the enzyme that disrupts the biofilm. This effect was specific to PelA_h_, as neither PslG_h_ nor the E165Q/E276Q variant were significant to ameliorate neutrophil killing, or affect neutrophil activity (**Fig 4B**). Combined, these data provide further evidence that PelA_h_ and PslG_h_ do not affect mammalian cell function, and that PelA_h_ can function to enhance neutrophil killing of *P. aeruginosa*.

**Enzymes effectively Disrupt Biofilms from Clinical and Environmental Isolates**. Our previous work established that clinical strains of *P. aeruginosa* can be divided into four different classes based upon their dependence on Pel and Psl exopolysaccharides for biofilm formation (*40*). Biofilm production in Class I strains is dependent on Pel, Class II strains are Psl dependent, Class III strains are redundant exopolysaccharide matrix producers and Class IV strains are matrix over-producers. In Class III and IV, both Pel and Psl contribute to biofilm formation. The effect of PelA_h_ and PslG_h_ on biofilm dispersal of isolates from each of the four classes was evaluated. Consistent with results previously reported, Class III environmental isolates exhibited poor attachment and biofilm formation while the cystic fibrosis clinical isolate CF127, a Class IV strain, produced the greatest biomass (*40*). Under the growth and assay conditions tested, treatment with 300 nM PslG_h_ +PelA_h_ for 2 h resulted in a 70–94% dispersal of biofilms formed by isolates from classes II-IV (**Fig 5**). PslG_h_ was more effective at disrupting biofilms from classes II-IV, suggesting that the Psl polysaccharide is a major contributor to biofilm biomass. An additive effect was observed for the matrix-overproducer CF127 where the combination of both enzymes led to the largest decrease in biofilm biomass (95% biomass reduction for PslG_h_+PelA_h_ compared to 85% for PslG_h_ alone). Strains PA14, the sole member of Class I and class IV member CF127 required 1 μM of enzyme to reduce the biofilm biomass by 58% and 95%, respectively. These data demonstrates that the enzymes are compatible with one another and can be utilized to disrupt biofilms of clinical and environmental *P. aeruginosa* isolates.

**Figure 5.**
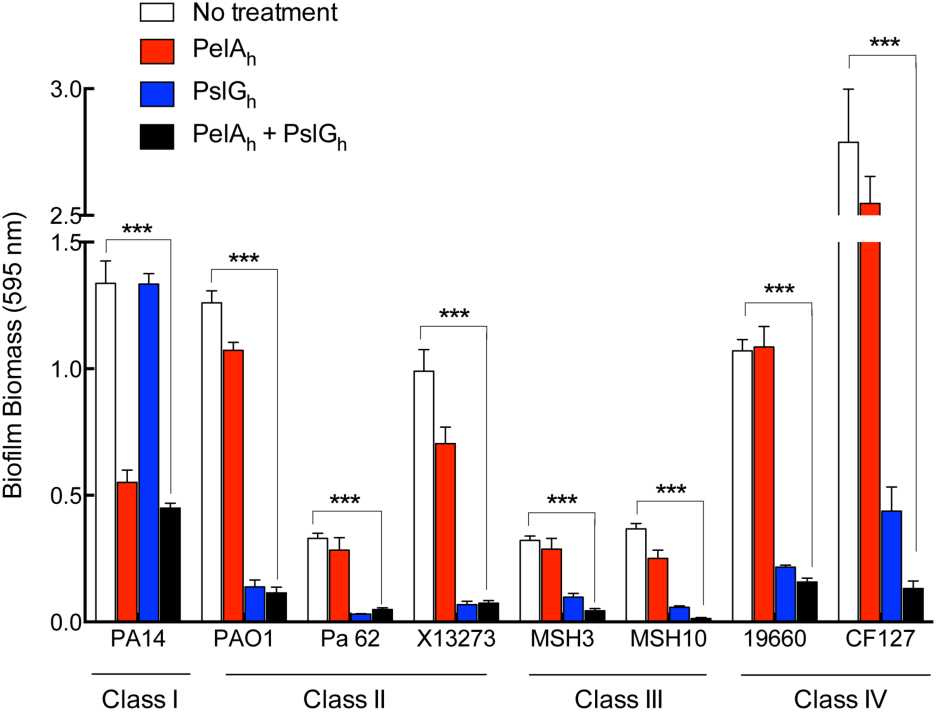
Enzymatic biofilm dispersal of P. aeruginosa clinical and environmental isolates. Isolates were grouped into categories as previously described and glycoside hydrolases PelA_h_ and PslG_h_ were exogenously added either individually or together and allowed to incubate for 2 h. Each data point represents the mean from three independent experiments of *n* = 3 crystal violet microtiter plate wells. Error bars indicate SEM. The *** indicates a significant difference (p ≤ 0.001).

## Discussion

In this work, we demonstrate that the glycoside hydrolases, PslG_h_ and PelA_h_, encoded in the exopolysaccharide biosynthetic operons of *P. aeruginosa* can be utilized to prevent and disrupt biofilms produced by the bacterium. We establish that these non-cytotoxic enzymes potentiate antibiotics and the innate immune system suggesting that they are promising enzymes for the treatment of *P. aeruginosa* infections. Since many Gram -positive and - negative bacterial exopolysaccharide biosynthetic operons encode a putative glycoside hydrolase or lyase, it is likely that this strategy could be extended to prevent and disrupt other exopolysaccharide-dependent biofilms. Examples include BcsZ, WssD, PssZ and PgaB involved in cellulose, acetylated cellulose, *Listeria monocytogenes* exopolysaccharide and poly-β-1,6-*N*-acetyl-D-glucosamine (PNAG) biosynthesis, respectively (*42, 43, 45, 46*). Glycoside hydrolase therapy therefore has the potential to target many Gram -positive and - negative biofilms from bacteria that are extremely relevant in healthcare and industrial settings.

Despite aggressive antimicrobial therapy, *P. aeruginosa* colonization is rarely eradicated and can lead to devastating biofilm-related persistent infections. Since exopolysaccharides that comprise the *P. aeruginosa* biofilm matrix aid in resistance to antimicrobial agents (7, *8, 39, 53*), studies have focused on inhibiting biofilm formation through the activation of bacterial response mechanisms. Halogenated furanones and their derivatives (*13–15*), cysteine sulfoxide compounds, (*16*), 6-gingerol (*17*) and meta-bromo-thiolactone (*18*) all target quorum-sensing signals to prevent *P. aeruginosa* biofilm formation. The use of Damino acids, which alter the cell envelope (*19*), and antimicrobial peptides (AMP) (*20–22*) which are capable of penetrating the biofilm have also been explored. Disappointingly, only a few molecules, including: nitric oxide (*54*), the fatty messenger *cis*-2-decenoic acid (CDA) (*55*) and the AMP 1080 (*22*), mediate both *P. aeruginosa* biofilm prevention and disruption. Nitric oxide decreases levels of the secondary messenger c-di-GMP levels, thereby down-regulating exopolysaccharide production and promoting planktonic growth (*56*), while CDA up-regulates genes involved in motility and metabolic activity and down-regulates genes contributing to attachment (*57*). Unfortunately, these molecules have only been tested on *P. aeruginosa* PAO1, and as the activity of these agents may not be specific to *P. aeruginosa*, they may negatively affect the natural microbiota. With the exception of *cis*-2-decenoic acid, these agents also require extended incubation times (≥ 24 h) to be efficacious against established biofilms. Therapeutic small molecules targeting cellular processes within the bacterial cytosol face numerous barriers including the biofilm matrix and two membranes that result in attrition. During this process many compounds are susceptible to multiple microbial resistance mechanisms including; impermeability of the membrane, sequestration, overproduction of the target, enzyme modification and multi-drug efflux.

Enzymes as therapeutics exhibit three inherent features that distinguish them from traditional small molecule drugs in that they bind and act on their targets with high affinity and specificity, their catalytic nature allows for the conversion of multiple target molecules, and because they target extracellular processes are prone to fewer resistance mechanisms. Therefore, the use of biofilm-dispersing enzymes has become recognized as a promising therapeutic strategy for the prevention and disruption of bacterial biofilms over the sole use of small molecule inhibitors, antibiotics and AMPs (*58–60*). DspB (DispersinB^®^), a glycoside hydrolase produced by *Actinobacillus actinomycetemcomitans*, hydrolyzes PNAG, a important exopolysaccharide required for biofilm formation and integrity by several Gram - positive and -negative pathogenic bacteria, but not *P. aeruginosa* (*47, 48, 61–64*). Dornase alfa (DNase I), which is in clinical use, disrupts bacterial biofilms through the hydrolysis of eDNA, hence reducing biofilm biomass and potentiating some antibiotics (*65, 66*). While young *P. aeruginosa* biofilms are sensitive to DNaseI treatment based on the role of eDNA during initial biofilm establishment, mature biofilms are less susceptible presumably since the exopolysaccharide changes during development (*67, 68*). Our results indicate that PslG_h_ and PelA_h_ are compatible with antibiotics and neutrophils, and function to rapidly and selectively target exopolysaccharide on the bacterial surface. This mechanism of action reduces susceptibility to host resistance mechanisms and off-target effects to the microbiota and host carbohydrates based upon the unique chemical structures of Pel and Psl. While the exopolysaccharide binds biofilm matrix components including eDNA and proteins to form a cohesive and structurally robust biofilm (*69, 70*), both glycoside hydrolases are able to access and hydrolyze the polysaccharides. Penetration and dispersal of the biofilm has several potential consequences including; increased penetration of antibiotics within the biofilm matrix, and the reduction of microenvironments that can render antimicrobials inactive. Since many infections are often polymicrobial, electing to target a single bacterial species may be perceived as inefficient. However, Pel has been demonstrated to promote mixed species biofilms with *Staphylococcus aureus* (*71*), and Psl-mediated protection against detergent stress and antibiotics is conferred to other pathogen bacteria in mixed species biofilms (*8, 72*). Therefore, hydrolysis of Pel and Psl in mixed species biofilms could increase the susceptibility of other embedded pathogenic bacteria to conventional therapeutic strategies.

In conclusion, bacterial biofilms are notoriously recalcitrant to immune defences and antibiotics and therefore novel therapeutic methods are urgently needed. Our study demonstrates that components of the *P. aeruginosa* exopolysaccharide biosynthetic operons can be manipulated to disrupt these biofilms allowing for antibiotic potentiation and effective killing by innate immunity. We hypothesize that glycoside hydrolases from other exopolysaccharide biosynthetic pathways can also be exploited to target diverse biofilms in a wide-range of applications.

## Materials and Methods

**Strains and culture conditions**. Strains used in this study are detailed in Table S1 and detailed culture conditions are described in *SI Materials and Methods*.

**Cloning, Expression, and Purification of PelA and PslG constructs**. PslG_h_ was purified as previously described (*49*). The DNA sequence of *pelA* from *P. aeruginosa* PAO1 was obtained from GenBank under accession no. AAG06452.1 (*73*). The PRED-TAT server (*74*) indicates that PelA possesses a TAT signal sequence from residues 1–45. Full experimental detail for cloning, expression and purification of PelA_h_ are provided in the *SI Materials and Methods*.

**Confocal Microscopy, Mitrotiter Dish Biofilm Assays and *P. aeruginosa* growth assays**.

Full experimental detail for confocal microscopy, mitrotiter dish biofilm assay and P. aeruginosa assays can be found in *SI Materials and Methods*.

**Antibiotic sensitivity assay**. Overnight cultures of *P. aeruginosa* PAO1 Δ*wspF* Δ*psl* P_BAD_*pel* (Pel-dependent) and *P. aeruginosa* PAO1 Δ*pelF* P_BAD_*psl* (Psl-dependent) were diluted to an OD_600_ of 0.05 in LB no salt (LBNS) with 0.5% arabinose and grown statically in polystyrene tubes in a final volume of 1 mL/tube. PelA_h_ and PslG_h_ were added at a final concentration of 2 μM and incubated with exopolysaccharide producing cells for 24 h at 25°C. Cultures which did not contain enzyme served as untreated samples. Following incubation, colistin was added to all tubes at a final concentration of 50 μg/mL and incubated for 24 h. Adherent biofilm and embedded cells were resuspended by scraping the tubes and vigorous pipetting to ensure removal of all cellular material and prevent experimental bias. Viability was quantified by serial dilutions and CFU counts on LB agar plates of the surviving population. Experiments were performed three times to obtain an average and standard error.

**Human neutrophil killing assay**. Overnight cultures of *P. aeruginosa* PAO1 Δ*wspF* Δ*psl* P_BAD_*pel* and *P. aeruginosa* PAO1 Δ*pelF* P_BAD_*psl* were diluted to an OD_600_ of 0.05 in LBNS with 0.5% L-arabinose and inoculated in a 96 well tissue culture-treated plate at a final volume of 100 μL/well. The plate was incubated statically at 28°C for 20 h. Supernatants were aspirated and 100 μL of phenol red-free Roswell Park Memorial Institute media (RPMI)+10% FBS containing 0.5 μM of PelA_h_ or PslG_h_ was added. The plate was incubated at room temperature on a Nutator for 1 h. Following pretreatment with hydrolase, 100 μL RPMI+10% FBS containing 6×10^6^ differentiated HL-60 cells were added to the wells, and plate was incubated for 90 min at 37 °C, 5% CO_2_. Wells were aspirated and supernatant was diluted between 1/200000 and 1/400000, and plated (50 μL) onto LB agar. To aspirated wells, 200 μL of 2 μM of PelA_h_ and 2 μM PslG_h_ was added and plate was incubated at room temperature on the Nutator for 1–1.5 h. Wells were aspirated, diluted and plated onto LB agar as above.

**Cell morphology and viability assays**. IMR-90 human lung fibroblast cells were seeded in 96-well CellBIND plates (Corning^®^) at a density of 8,000–10,000 cells/well. The next day, the media was exchanged with media containing 1 μM CellTracker Orange CMRA (Molecular Probes) in serum-free EMEM (Wisent) media or supplemented with 10% FBS. After 60 min, excess dye was removed by media exchange. PslG_h_ and PelA_h_ were added to a final concentration of 1 mg/mL. The detergent digitonin and the *Clostridium difficile* toxin TcdB were utilized as controls and added to a final concentration of 0.03 mg/mL and 0.5 pM, respectively (*51*). The cell plates were returned to the incubator for 5.0 h before imaging. CellTracker-labeled cells were evaluated on a Cellomics ArrayScan VTI HCS reader (Thermo Scientific) using the target acquisition mode, a 10x objective, and a sample rate of 100 objects per well. Following 24 h incubation, cells in serum-free media were supplemented with 10% FBS. At 48 h post incubation, PrestoBlue^®^ reagent was added at a 1:10 reagent:media to the cells and allowed to incubate for 5.0 h. The microtiter plates were read in a SpectroMax M2 plate reader with a λ_x_ of 555 nm and λ_em_ of 585 nm. Western blotting to probe for the presence of PslG_h_ and PelA_h_ was completed using polyclonal antibodies that were previous generated against both enzymes (*33, 49*).

**Statistical Analysis**. One-way ANOVA and Tukey’s multiple comparison test were utilized to valuate statistical significance. EC_50_ values were calculated using non-linear least-squares fitting to a dose-response model in Prism (GraphPad Software) All means, SEM, bar graphs and dose-response curves were calculated and generated using Prism 6.0.

## ACKNOWLEDGEMENTS

^*^Research described in this paper is supported by operating grants from the *Canadian Institutes of Health Research* (CIHR) (#43998 to P.L.H., #123306 to D.C.S., #286650 to R.A.M., and #81361 to P.L.H. and D.C.S.), *Cystic Fibrosis Canada* (CFC) (D.C.S. and P.L.H.), the *National Institutes of Health* (R01AI097511 to D.J.W. and 2R01AI077628 to M.R.P.) and the Natural Sciences and Engineering Research Council of Canada (RGPIN 418405 to R.A.M.). P.B. has been supported in part by a CFC postdoctoral fellowship and a Banting Fellowship from CIHR. B.D.S has been supported by graduate scholarships from CFC and CIHR. L.K.J. is the recipient of an American Heart Association Postdoctoral Fellowship (14POST20130017). P.L.H is the recipient of a Canada Research Chair.

## Supplementary Material and Methods

### Cloning, Expression, and Purification of PelA_h_ and PslG_h_ constructs

To obtain a soluble protein constructs, *pslG* and *pelA* were amplified from genomic DNA by PCR using the primers CT<u>GCATATG</u>GGCGGGCCGTCCAGCGTGGCG and TTT<u>CTCGAG</u>TCACGGTTGCAC CTCGACGTC, respectively. Introduced *Nde*I, and *Xho*I restriction sites are underlined and each gene was ligated into the pET28a (Novagen) expression vector encoding an N-terminal His-tag. This generated PelA_47–303_. All site-directed mutagenesis to generate protein variants was performed using the QuikChange^®^ Lightning kit according to the prescribed protocol (Agilent Technologies). Constructs generated were verified by sequencing performed by ACGT DNA Technologies Corporation (Toronto, ON).

*E. coli* BL21 (DE3) CodonPlus cells (Stratagene) were transformed with the expression plasmid and grown in 2 L Lauria-Bertani (LB) broth containing 50 μg/mL kanamycin at 37 °C. When the OD_600_ of the cell culture reached 0.5–0.6, protein expression was induced by the addition of isopropyl β-D-1-thiogalactopyranoside (IPTG) to a final concentration of 0.5 mM. The cells were incubated post-induction overnight at 18 °C with shaking at 200 rpm before being harvested by centrifugation at 5,000 × g for 30 min at 4 °C. Cell pellets containing PelA_47–303_ were resuspended in 40 mL of buffer A (20 mM imidazole, 50 mM Tris-HCl pH 7.5, 300 mM NaCl, 2% (v/v) glycerol) plus one SIGMA*FAST*™ Protease Inhibitor Tablet. The cells were lysed by at least three passes through an Emulsiflex C3 homogenizer at 100 MPa (Avestin Inc) and the resulting cell debris was separated from soluble protein by centrifugation at 35,000 × *g* for 30 min. The supernatant was applied to 5 mL of Ni-NTA Superflow resin packed into a gravity column (Qiagen) pre-equilibrated with buffer A. The column was washed with 3 column volumes (CV) of buffer A and the expressed protein was eluted with buffer A1 supplemented with 250 mM imidazole. The eluted fractions were concentrated to a 1–2 mL volume using an Amicon Ultra centrifugation filter device (Milipore) with a 10 kDa cutoff and the protein was further purified via size-exclusion chromatography using a HiLoad 16/60 Superdex 200 gel-filtration column (GE Healthcare). The purity of protein was judged to be >95% pure by SDS–PAGE and the protein could be concentrated to 8–10 mg/mL and stored at 4 °C for at least one month without precipitation or degradation.

**Confocal Microscopy**. Psl biofilms were grown overnight at room temperature in uncoated 15 μ-Slide VI^0.4^ flow cell chambers (ibidi GmbH, Martinsfried, Germany). The channels were inoculated with 200 μL of a culture with an OD_600_ of 0.5 grown in LBNS supplemented with 0.5% arabinose. Biofilms were washed three times with sterile PBS and then treated with PslG_h_, PslG_h_ E165Q/E276Q and buffer only control [50 mM Tris pH 7.5, 150 mM NaCl, 10% (v/v) glycerol] statically for 1 h at room temperature. The final enzyme concentration was 86 nM. After digestion, biofilms were stained with FITC conjugated *Hippeastrum Hybrid Amaryllis* (HHA; EY Labs, San Mateo, CA) lectin at 100 mg/mL for two hours at 4 °C as previously described ^1^. The biofilms were then washed and fixed with 4% paraformaldehyde. Fluorescent images were acquired using an Olympus FV1000 Filter confocal system using a 20x LUCPLFLN, N.A. 0.45 objective lens (Olympus America Inc, Melville NY). Images were analyzed and constructed using Olympus Fluoview version 03.01 software.

Pel-dependent biofilms were cultivated as described previously with minor modifications ^2^. Flow cell chambers were inoculated with a mid-log LB culture of *P. aeruginosa* PA14 that was diluted with glucose minimal media (0.3 mM final glucose concentration) to an OD_600_ of 0.01. Cells were allowed to attach for 1 h before induction of flow. Biofilms were grown on glucose minimal media for two days at room temperature at a constant flow rate (10 mL/h) before treatment with PelA_h_, PelA_h_ E218A, and buffer only control [20 mM Tris pH 8.0, 150 mM NaCl, 10% (v/v) glycerol] statically for 1 h at room temperature. The final enzyme concentration was 85 nM. After digestion, biofilms were washed; and then, Pel was stained with fluorescein-labeled *Wisteria floribunda* lectin (WFL, 100 μg/mL, Vector Laboratories) for 15 minutes. Stained biofilms were washed prior to visualization on a Zeiss LSM 510 scanning confocal laser microscope. Image analysis was conducted using Velocity software (Improvision). Experiments were performed in biological duplicate.

**Microtiter dish biofilm assay**. To examine biofilm prevention, Psl- and Pel-arabinose inducible *P. aeruginosa* PAO1 (PAO1 Δ*pelF* P_BAD_*psl* and PAO1 Δ*wspF* Δ*psl* P_BAD_*pel*) were grown at 37 °C overnight with shaking at 200 rpm. The cultures were normalized to an OD_600_ of 0.5 and then diluted 1:100 in LB without salt (LBNS). L-arabinose was added to a final concentration of 0.5% (w/v) to induce exopolysaccharide biosynthesis and biofilm formation. 95 μL of diluted culture was added to sterile 96-well polystyrene microtiter plates (Thermo Scientific Cat No. 243656) and varying concentrations of PelA or PslG (0.1 nM – 5 μM) were added in 5 μL aliquots to give a final volume of 100 μL. The cultures were incubated statically for 24 h at 25 °C to allow for biofilm formation. To eliminate edge-effects, ~200 μL of sterile water was placed in all outside wells and the plate was sealed with parafilm. After incubation, non-adherent cells and media were removed by thoroughly washing the plate with deionized water. The wells were stained with 150 μL of 0.1% (w/v) crystal violet for 10 min followed by rinsing with water. The remaining dye was solubilized by addition of 150 μL of 95% (v/v) ethanol and left for 10 min after which time the absorbance was measured at 595 nm using a SpectraMax M2 from Molecular Devices (Sunnyvale, CA). The amount of biofilm is proportional to the absorbance from staining with crystal violet ^3^.

For biofilm disruption assays, biofilm cultures were grown statically for 24 h. Following incubation, non-adherent cells and media were removed by washing the plate with dH_2_O. The wells were filled with 95 μL of 100 mM sodium HEPES buffer pH 7.0 followed by 5 μL of varying concentrations of each hydrolytic enzyme (2 nM – 5 μM). Reactions were allowed to proceed for up to 60 min at 25 °C on a rotating nutator at which time, the reaction was quenched by washing the plates with dH_2_O. The wells were stained with 150 μL of 0.1% (w/v) crystal violet for 10 min, washed and solubilized with ethanol prior to quantification. All reactions were completed in triplicates. The addition of 2.5 mg/mL of kanamycin to culture prior to biofilm formation was used as positive control.

***P. aeruginosa* growth assay**. To assay for glycoside hydrolase cytotoxicity to P. aeruginosa PAO1, a bacterial growth assay was set up as described for the biofilm inhibition assay with the addition of 25 μM of PelA_h_ or PslG_h_ added at the time of inoculation. The bacteria were grown statically at 37 °C in a thermo-controlled SpectraMax M2 spectrophotometery. At 30 min intervals, the OD_600_ of each culture was measured for a duration of 6 h. An untreated culture was used as a control.

## FIGURE LEGEND

**Figure S1.**
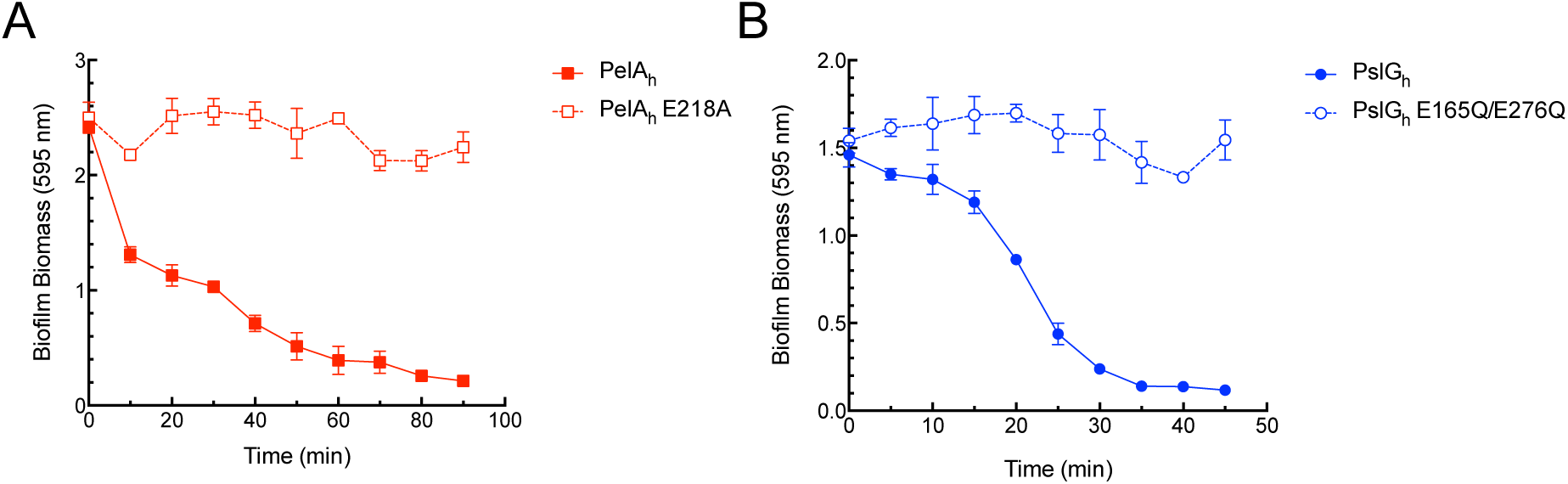
Time course disruption of *P. aeruginosa* biofilms. Crystal violet staining of biofilms following the exogenous addition of glycoside hydrolases (A) PelA_h_ and (B) PslG_h_ on their respective exopolysaccharide. Each data point represents the mean from *n* = 3 crystal violet microtiter plate wells. Error bars indicate SEM.

**Figure S2.**
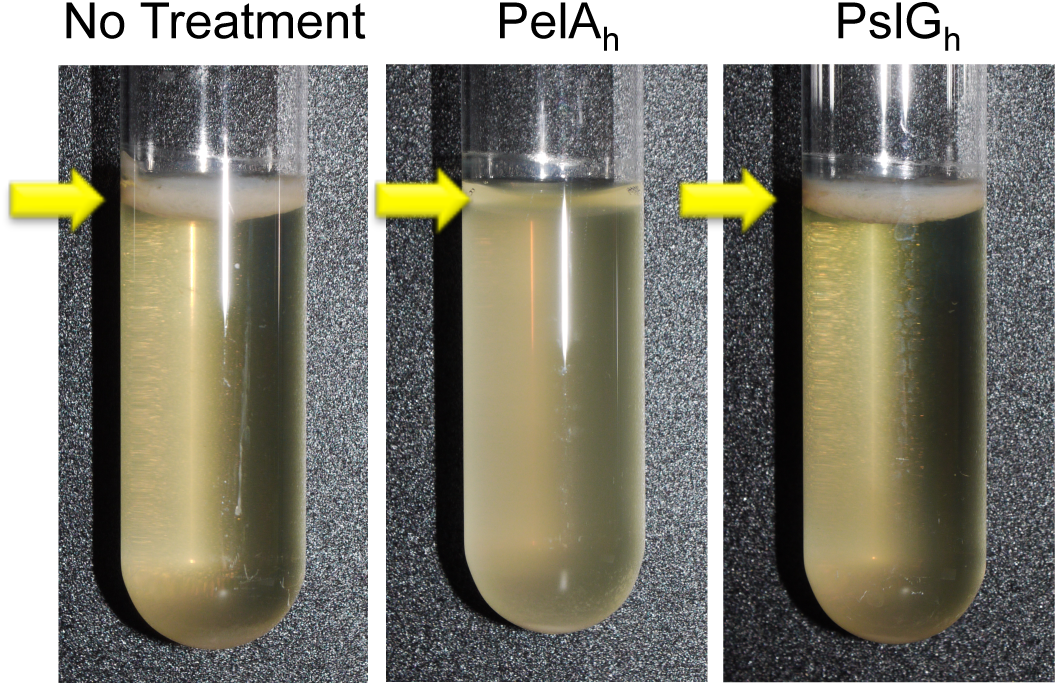
Biofilm prevention standing culture pellicle assay. Biofilm formation at the air-liquid interface was examined in Pel-dependent culture following incubation with PelA_h_ and PslG_h_. Arrows indicate the location of the air-liquid interface where biofilm formation occurs.

**Figure S3.**
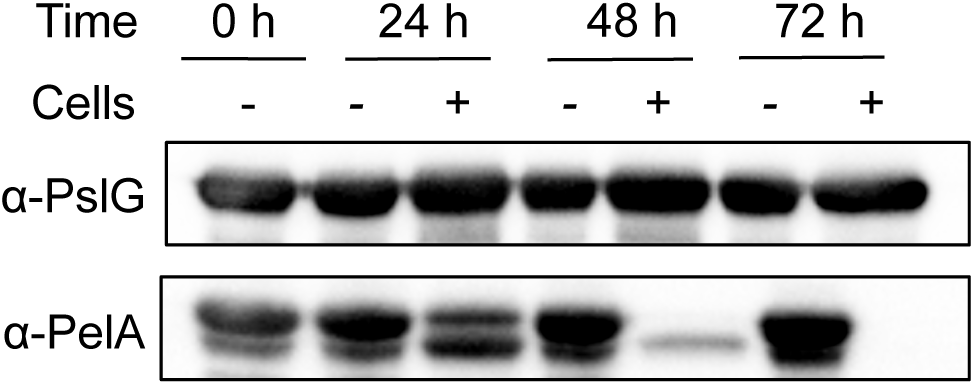
Protein stability of PelA_h_ and PslG_h_ in *P. aeruginosa* culture. Western blotting using *α*-PelA and α-PslG to detect the presence of exogenously applied PelA_h_ and PslG_h_ at various time points during incubating with P. aeruginosa Pel and Psl biofilm formation, respectively. Incubation of each glycoside hydrolase in the absence of *P. aeruginosa* culture (Cell -) was utilized for comparison.

**Figure S4.**
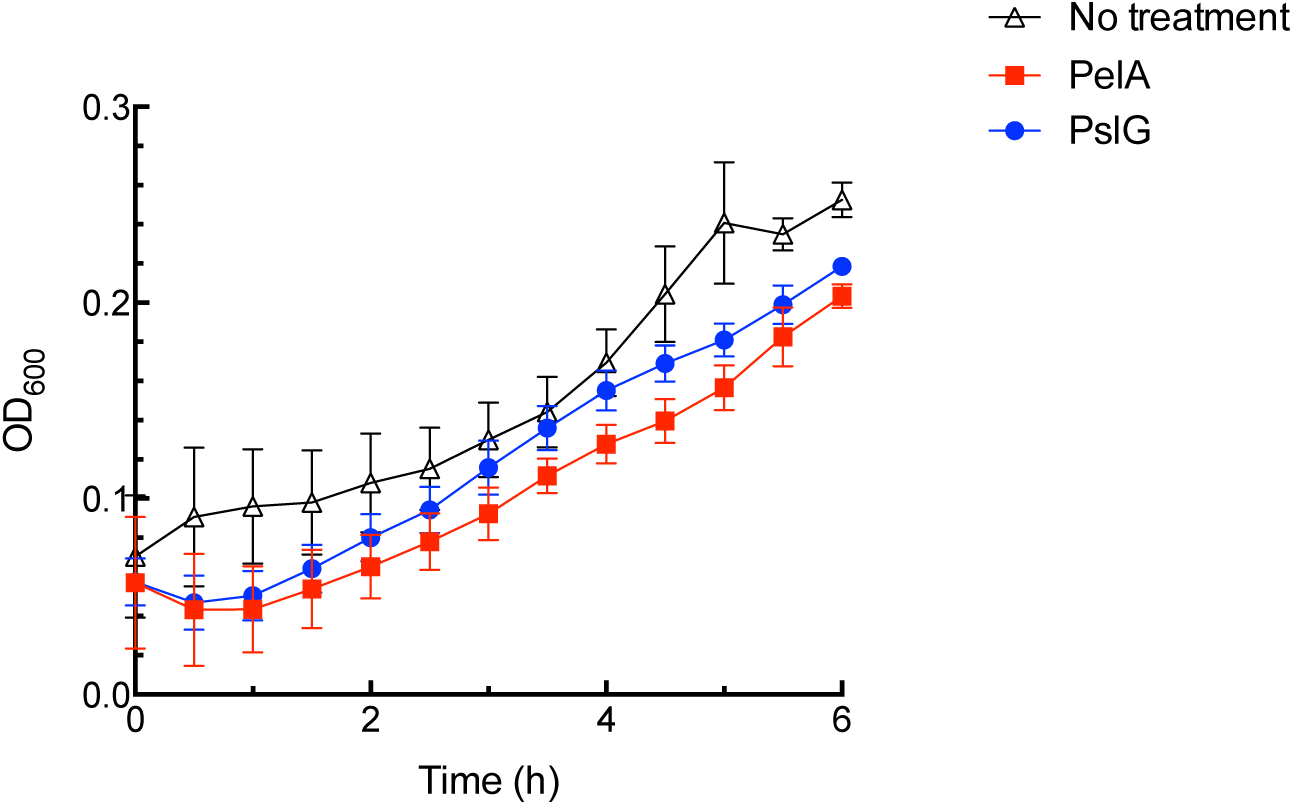
Glycoside hydrolases do not affect *P. aeruginosa* growth. Growth curve with *P. aeruginosa* PAO1 in the presence of PslG_h_ and PelA_h_ over 6 h at 37 °C.

**Figure S5.**
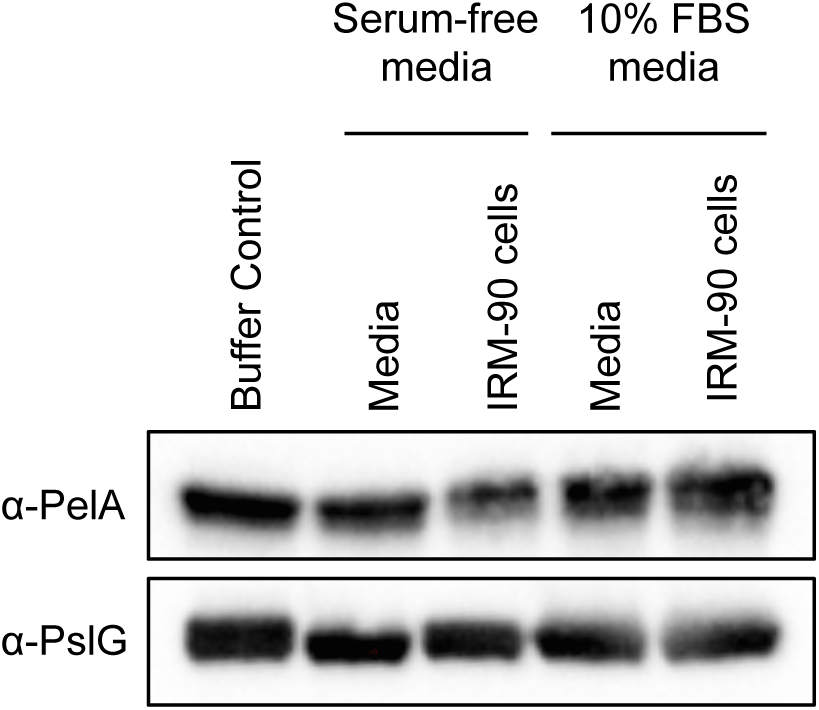
Protein stability of PelA_h_ and PslG_h_ in mammalian cell culture. Western blotting of exogenously added PelA_h_ and PslG_h_ after 48 h incubation in IMR-90 cell culture in the presence and absence of 10% FBS in the media.

**Table S1:**
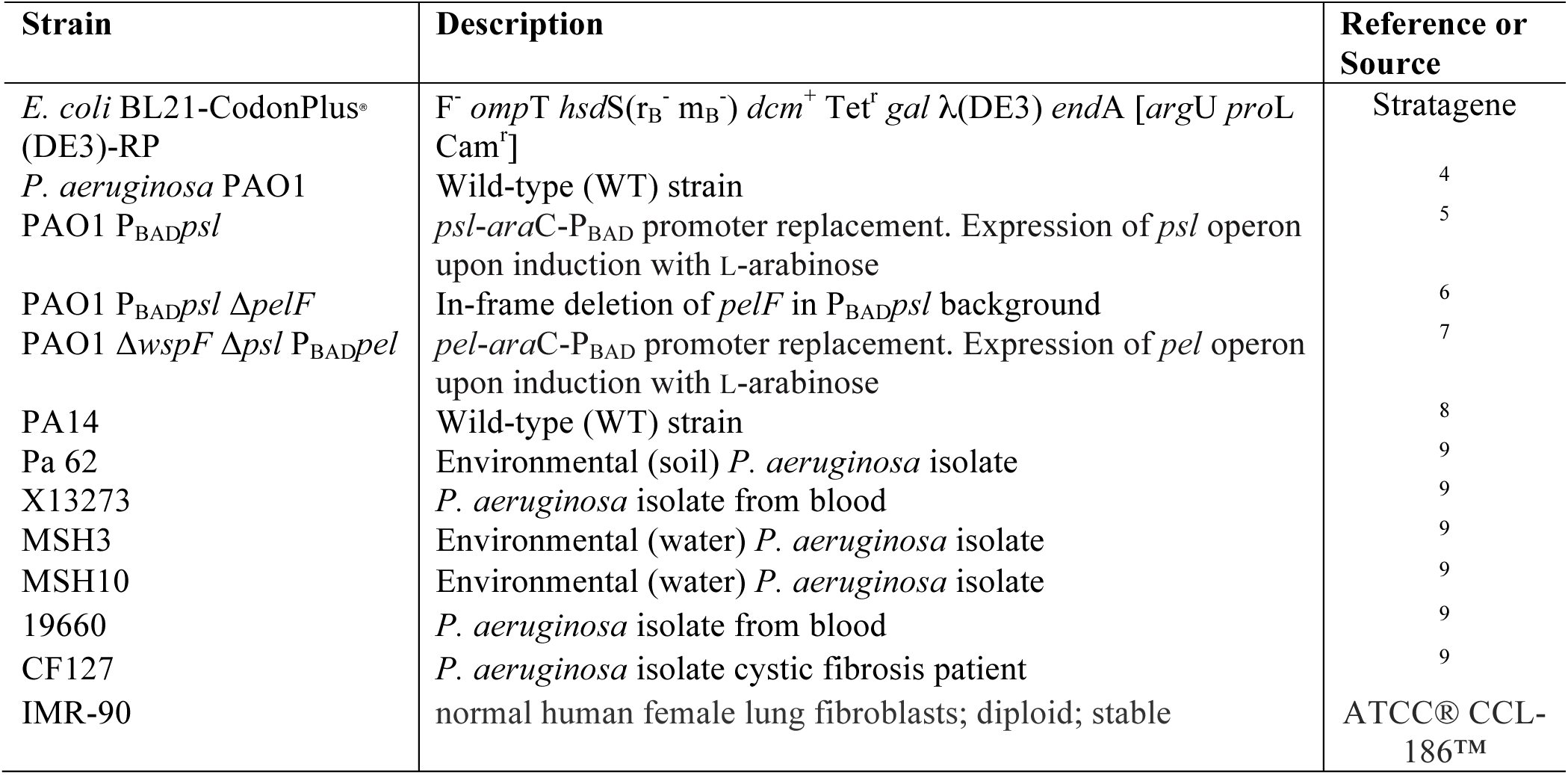
Strains and Plasmids used in this Study.

## Footnotes

* Research described in this paper is supported by operating grants from the Canadian Institutes of Health Research (CIHR) (#43998 to P.L.H., #123306 to D.C.S., #286650 to R.A.M., and #81361 to P.L.H. and D.C.S.), Cystic Fibrosis Canada (CFC) (D.C.S. and P.L.H.), the National Institutes of Health (R01AI097511 to D.J.W. and 2R01AI077628 to M.R.P.) and the Natural Sciences and Engineering Research Council of Canada (RGPIN 418405 to R.A.M.). P.B. has been supported in part by a CFC postdoctoral fellowship and a Banting Fellowship from CIHR. B.D.S has been supported by graduate scholarships from CFC and CIHR. L.K.J. is the recipient of an American Heart Association Postdoctoral Fellowship (14POST20130017). P.L.H is the recipient of a Canada Research Chair.

